# Sex and race shape serum protein biomarkers and bias diagnostic reference intervals

**DOI:** 10.64898/2026.07.18.739384

**Authors:** Ahrum Son, Jaeho Ji, Eunjeong Han, Yunho Choi, Jumin Park, Heejin Lee, Sujin Park, Hyunsoo Kim

## Abstract

Clinical protein biomarkers—whether measured by FDA-cleared in-vitro diagnostic (IVD) assays or deployed as laboratory-developed tests (LDTs)—are routinely interpreted against fixed, population-agnostic reference intervals. Whether the baseline serum concentrations of these proteins differ systematically by sex and by self-reported race in healthy individuals has not been tested across a broad clinical panel on a single standardized platform. Using scheduled multiple reaction monitoring – mass spectrometry (MRM-MS) with stable-isotope-labelled internal standards, we quantified 87 proteins (39 FDA-cleared analytes; 48 LDT/research markers) in 60 healthy, age-matched donors balanced across three races (White, Hispanic, Black) and both sexes. Sex was a pervasive axis of variation: donors separated by sex in unsupervised multivariate space (PERMANOVA p = 0.020) but not by race (p = 0.14). Three FDA-target proteins differed by sex at FDR q < 0.05—sex hormone-binding globulin, haemoglobin-α and albumin—with directions preserved within every race. Race differences were real but protein-specific, exemplified by ancestry-associated CD14. Critically, applying a single pooled 95% reference interval mis-flagged as many as ∼25% of a demographic subgroup as abnormal, versus the 5% expected. Demographic context should therefore be incorporated into reference intervals and decision thresholds for protein-based diagnostics.

## Introduction

Protein biomarkers underpin a large share of modern laboratory medicine. Some are the analytes of assays cleared or approved by the U.S. Food and Drug Administration (FDA) as in-vitro diagnostics (IVDs); many more are deployed as laboratory-developed tests (LDTs) or remain research-grade markers under active clinical evaluation. The distinction is regulatory rather than biological, yet it has become newly consequential, as the shifting oversight landscape for LDTs has intensified scrutiny of how such tests are validated and interpreted before they inform patient care.^1^ Whatever their regulatory status, however, these assays share a common interpretive logic that has changed remarkably little in decades.

In routine practice, a measured concentration is compared against a reference interval or a fixed decision threshold, and the patient is classified as normal or abnormal accordingly. The validity of that single act of classification rests on an implicit and rarely examined assumption—that the reference interval derived from some reference population actually applies to the individual being tested.^2^ Reference intervals are, by construction, statistical summaries of a “healthy” population, and international guidance has long recognized that they may require partitioning when a physiological covariate shifts the underlying distribution.^3^ In practice, though, many protein assays are still read against a single, pooled interval applied uniformly across sex and ancestry.

Physiology gives ample reason to doubt that a one-size-fits-all interval holds uniformly. Sex is among the most pervasive axes of biological variation in the human body, shaping hormonal milieu, body composition, haematological set-points and hepatic synthetic capacity, and it leaves a broad imprint across the genome and transcriptome.^4^ The recognition that sex operates as a fundamental biological variable—rather than a nuisance to be averaged away—now underpins research policy and a growing appreciation that diagnostic and therapeutic responses can differ systematically between women and men.^5, 6^

Ancestry adds a second, more contested layer. A substantial literature documents ancestry-associated differences in the baseline levels of specific plasma proteins, some traceable to allele-frequency differences at the encoding locus—soluble CD14 being a textbook example.^7^ At the same time, the uncritical insertion of race into clinical algorithms has come under sustained re-examination, because self-reported race is a social construct that captures genetic ancestry only imperfectly and can encode bias when treated as a biological variable.^8, 9^ The removal of race coefficients from estimated glomerular filtration rate and from pulmonary-function equations has demonstrated both the clinical weight of these adjustments and the hazards of conflating race with biology.^10, 11, 12, 13, 14^ Distinguishing genuine, mechanistically grounded biological differences from spurious or socially mediated ones is therefore essential—and doing so requires measurement, not assumption.^15, 16^

Large-scale plasma proteomics has begun to map this variation at breadth. Affinity-based surveys of tens of thousands of participants report that age, sex and genetics jointly shape thousands of circulating proteins, and that these signatures carry disease-predictive information.^17, 18, 19, 20, 21^ These efforts are transformative, yet most rely on multiplexed affinity platforms whose readouts are relative and platform-specific, and most draw on cohorts of predominantly European ancestry. Crucially, when evidence is assembled piecemeal—one protein, one assay, one cohort at a time—true biological differences become entangled with cross-assay analytical variability, and the demographic signal that matters for diagnosis is difficult to isolate.^22^

Targeted mass spectrometry offers a complementary and, for this question, uniquely well-suited approach. Scheduled multiple-reaction-monitoring (MRM) with stable-isotope-labelled standard (SIS) peptides measures many pre-specified proteins simultaneously against an internal reference, yielding quantitation that is specific, reproducible and portable across laboratories and over time.^23, 24, 25, 26, 27, 28^ These are precisely the properties needed to compare demographic subgroups on a common analytical footing, and they are why MRM has become a reference method for bridging biomarker discovery to clinical verification.

Here we exploit that footing to address the question directly. Using a single scheduled-MRM assay, we measured 87 clinically used biomarker proteins—spanning FDA-cleared IVD analytes and LDT/research markers—in 60 healthy, age-matched donors balanced by race and sex. Our central question is deliberately practical: do the baseline serum concentrations of these proteins differ by race and by sex in healthy people, and if so, by enough to matter when the same proteins are used to diagnose disease? We find that sex is a pervasive, direction-consistent axis of variation, that race differences are real but concentrated in specific proteins, and that ignoring either when setting reference intervals produces a systematic, quantifiable misclassification of healthy individuals.

## Material & Methods

### Cohort and serum samples

Sixty commercially sourced (Innovative Research Inc, Novi, MI, USA) single-donor human serum were analyzed, balanced into six race–sex groups of ten donors each: White (Caucasian), Hispanic and Black (African American), each split evenly by sex. All donors were healthy young adults (age range 20–30 years; overall mean 25.0 ± 3.4 years). Age distributions were comparable across the six groups (group means 24.4–25.6 years), removing age as a confounder of the demographic contrasts. Donor identifiers follow a {race}{sex}_{number} scheme (e.g. WM_001 = White male donor 1). Per-donor age, sex, race and manufacturing-lot metadata are provided in **Table S1** and summarized in **Table 1**.

**Table 1.**

| Race | Sex | n | Age mean±SD | Age range |
| --- | --- | --- | --- | --- |
| White | Male | 10 | 25.6±3.9 | 20~30 |
| White | Female | 10 | 24.6±3.1 | 20~29 |
| Hispanic | Male | 10 | 24.5±3.6 | 20~30 |
| Hispanic | Female | 10 | 25.3±3.7 | 20~29 |
| Black | Male | 10 | 24.4±3.3 | 20~29 |
| Black | Female | 10 | 25.6±3.3 | 21~30 |
| All | Both | 60 | 25.0±3.4 | 20~30 |

### Protein digestion

Serum samples were prepared for MRM-MS-based quantitative proteomic analysis by enzymatic digestion. Each tube, containing 100 μg of protein, was first treated to reduce disulfide bonds using 2% sodium deoxycholate (SDC) and 10 mM tris-(2-carboxyethyl)-phosphine hydrochloride (TCEP) at 60□ for 60 minutes. To block free cysteines, 20 mM iodoacetamide (IAA) was added and the samples were incubated in the dark at 25□ for 30 minutes. The protein solution was then diluted to a final concentration of 20 mM ammonium bicarbonate (ABC) with distilled water. Trypsin (Promega, Sequencing Grade Modified, 20 μg) was added at a 1:50 enzyme-to-protein ratio and digestion proceeded overnight at 37□. The reaction was quenched with 1% formic acid.

### MRM-MS assay and quantitation

Each serum was subjected to tryptic digestion and spiked with stable-isotope-labelled standard (SIS) peptides corresponding to the endogenous target peptides. Scheduled multiple-reaction-monitoring (MRM) LC-MS/MS acquisition monitored 93 peptides mapping to 87 proteins, with three fragmention transitions per peptide and paired light (endogenous) and heavy (SIS) channels. The peak-area ratio (PAR) — the primary quantitative readout — was computed for each peptide in each donor as:

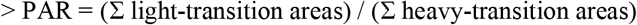

This transition-summed formulation weights each transition by its intensity and is more robust than the mean of per-transition ratios when individual transitions are near the noise floor (median inter-transition CV ≈ 41%; only ∼35% of transitions below 20% CV). Where a protein was represented by more than one peptide, the protein-level value was taken as the median of its peptides. All 93 peptides were quantified in all 60 donors (no missing values), spanning approximately six orders of magnitude in PAR.

### FDA & LDT target annotation

Each of the 87 proteins was manually annotated (**Table S1**) as either the analyte of an FDA-cleared/approved in-vitro diagnostic (IVD) assay (“FDA”; n = 39) or a laboratory-developed-test /research biomarker (“LDT”; n = 48), and assigned to one of eleven broad clinical categories (Coagulation/Platelet, Oncology, Complement, Inflammation/Immune, Endocrine/Metabolic, Lipid/Cardiovascular, Liver/Nutrition, Iron/Hematology, Renal, ECM/Other, Other). The FDA/LDT designation reflects the clinical status of an assay for that analyte and is intended to indicate the regulatory context in which the protein is used, not a claim about any specific commercial product.

### Statistical analysis

Protein-level PAR values were transformed as log□(PAR + m/2), where m is the smallest positive value in the matrix. Each protein was modelled by two-way analysis of variance:

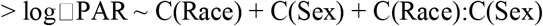

using type-II sums of squares. Partial η^2^ was computed per effect as the effect sum of squares divided by (effect + residual) sums of squares. Raw p-values were adjusted across the 87 proteins separately for each of the three effects using the Benjamini–Hochberg false-discovery-rate (FDR) procedure; adjusted values are reported as q. Pairwise differences between the three races were evaluated by Tukey’s honestly-significant-difference test. The equivalent peptide-level ANOVA (93 peptides) gave concordant conclusions.

Multivariate structure was assessed by principal-component analysis (PCA) on z-scored log□ values. Group separation was tested by permutational multivariate ANOVA (PERMANOVA) on Euclidean distances with 9,999 permutations, computed separately for race and for sex.

### Reference-interval misclassification simulation

For each analyzed FDA-target protein, a pooled 95% reference interval was defined on the log scale as (pooled mean ± 1.96 × pooled SD), using all 60 donors. For each of the six race–sex subgroups, the expected proportion of individuals falling below or above these pooled limits was computed analytically from the subgroup’s fitted normal (log-scale) distribution — P(x < lower) + P(x > upper). A perfectly calibrated, demographically unbiased reference interval yields 5% outside for every healthy subgroup; values above 5% quantify systematic over-flagging, values below 5% quantify under-flagging. The parametric formulation avoids the quantization artefacts of empirical counts at n = 10 per group. Per-subgroup, per-protein values are tabulated in **Table S4**.

## Results

### A single-platform panel of 87 clinical-target proteins across a balanced cohort

We assembled a cohort of 60 commercially sourced single-donor sera, balanced into six race–sex groups of 10 donors each (White, Hispanic and Black; male and female; **Table 1, Figure 1a–c**). All donors were healthy young adults (age 20–30 y; mean 25.0 y), with age distributions closely matched across groups, so that demographic contrasts are not confounded by age or by overt disease. Each serum was analyzed by scheduled MRM, yielding relative quantitation of 93 peptides mapping to 87 proteins. Of these, 39 are analytes of FDA-cleared IVD assays and 48 are used as LDT or research biomarkers; together the panel spans coagulation and platelet biology, oncology, complement, inflammation and immunity, endocrine and metabolic function, lipid and cardiovascular markers, liver and nutrition, iron and haematology, and renal categories (**Figure 1d, Table S1**). This breadth was deliberate: it lets us ask whether demographic effects are idiosyncratic to a few proteins or a general property of the clinically used proteome.

**Figure 1.**
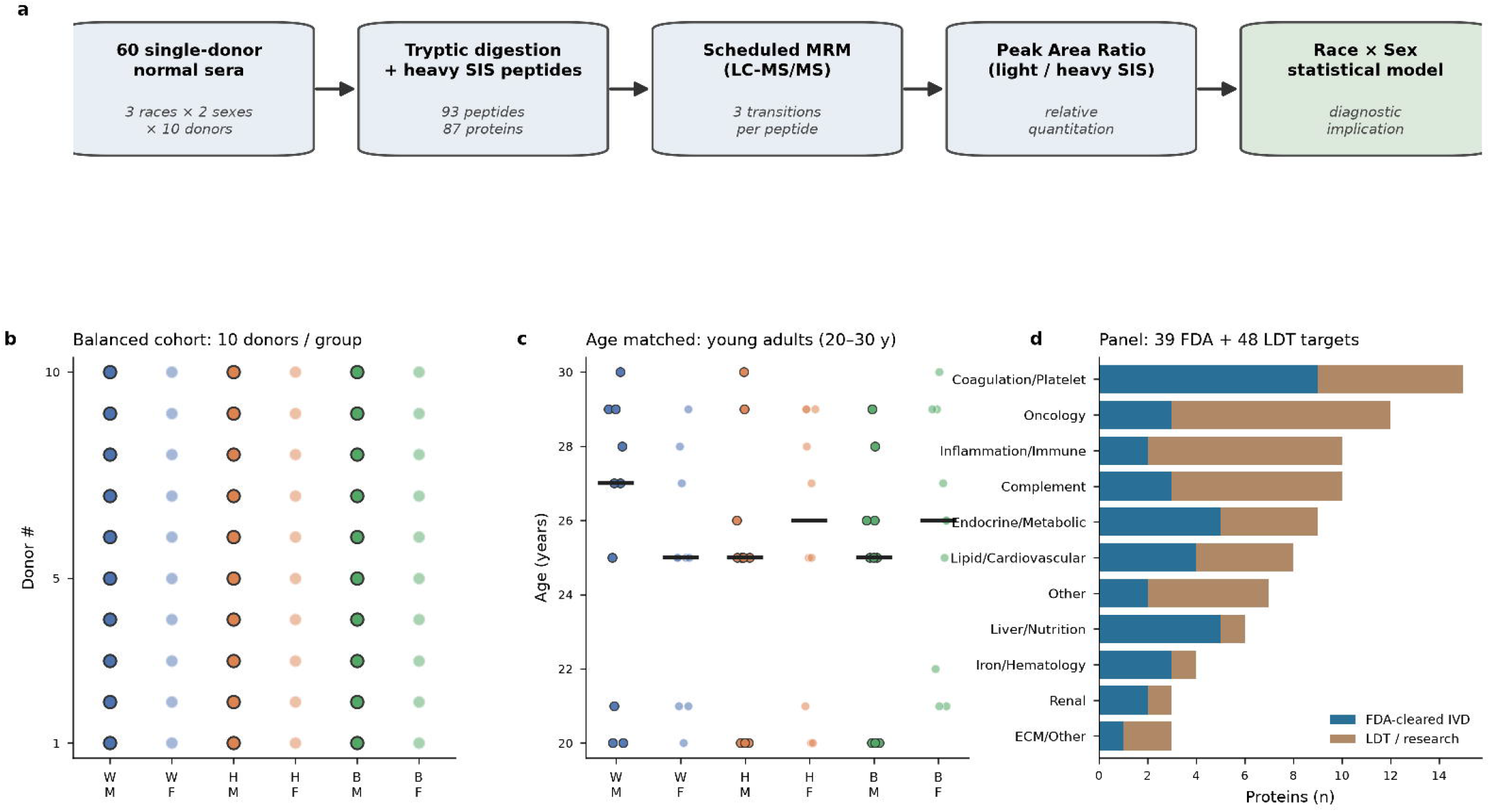
Study design: race- and sex-stratified serum MRM proteomics of clinical biomarker targets. (a) Analytical workflow: 60 single-donor normal sera (3 races × 2 sexes × 10 donors) underwent tryptic digestion with spiked stable-isotope-standard (SIS) peptides (93 peptides, 87 proteins), scheduled multiple-reaction-monitoring (MRM) LC-MS/MS with three transitions per peptide, peak-area-ratio (light/heavy SIS) relative quantitation, and race × sex statistical modelling with diagnostic interpretation. (b) Balanced cohort of 10 donors per race–sex group (WM, WF, HM, HF, BM, BF); dark outline denotes male and light fill denotes female. (c) Age-matched young-adult donors (20–30 y); horizontal bars indicate group medians. (d) Composition of the 87-protein panel by clinical category, coloured by regulatory status (FDA-cleared IVD versus LDT/research); the panel comprises 39 FDA and 48 LDT targets.

Assay performance supported quantitative comparison across this range. All 93 peptides were detected in all 60 donors, spanning roughly six orders of magnitude in abundance, with a median inter-transition coefficient of variation of ∼41% (**Figure S1**). Peptide-level values were computed as the transition-summed light/heavy ratio—more robust than the mean of per-transition ratios when individual transitions are noisy—and aggregated to the protein level by the median of constituent peptides. The internal SIS reference for every peptide ensures that the between-donor differences we report reflect endogenous abundance rather than run-to-run analytical drift.

### Sex is a systematic axis of variation; race is protein-specific

We first examined global structure without reference to any single protein. In principal-component space, donors did not separate by race (PERMANOVA F = 1.21, p = 0.14; **Figure 2a**), whereas they did separate by sex (F = 1.83, p = 0.020; **Figure 2b**). Unsupervised hierarchical clustering of the most variable proteins was organized by overall abundance rather than by any race or sex block (**Figure 2c**; full-panel clustering in **Figure S2**), indicating that sex acts as a distributed signal spread across many proteins rather than one carried by a few dominant markers.

**Figure 2.**
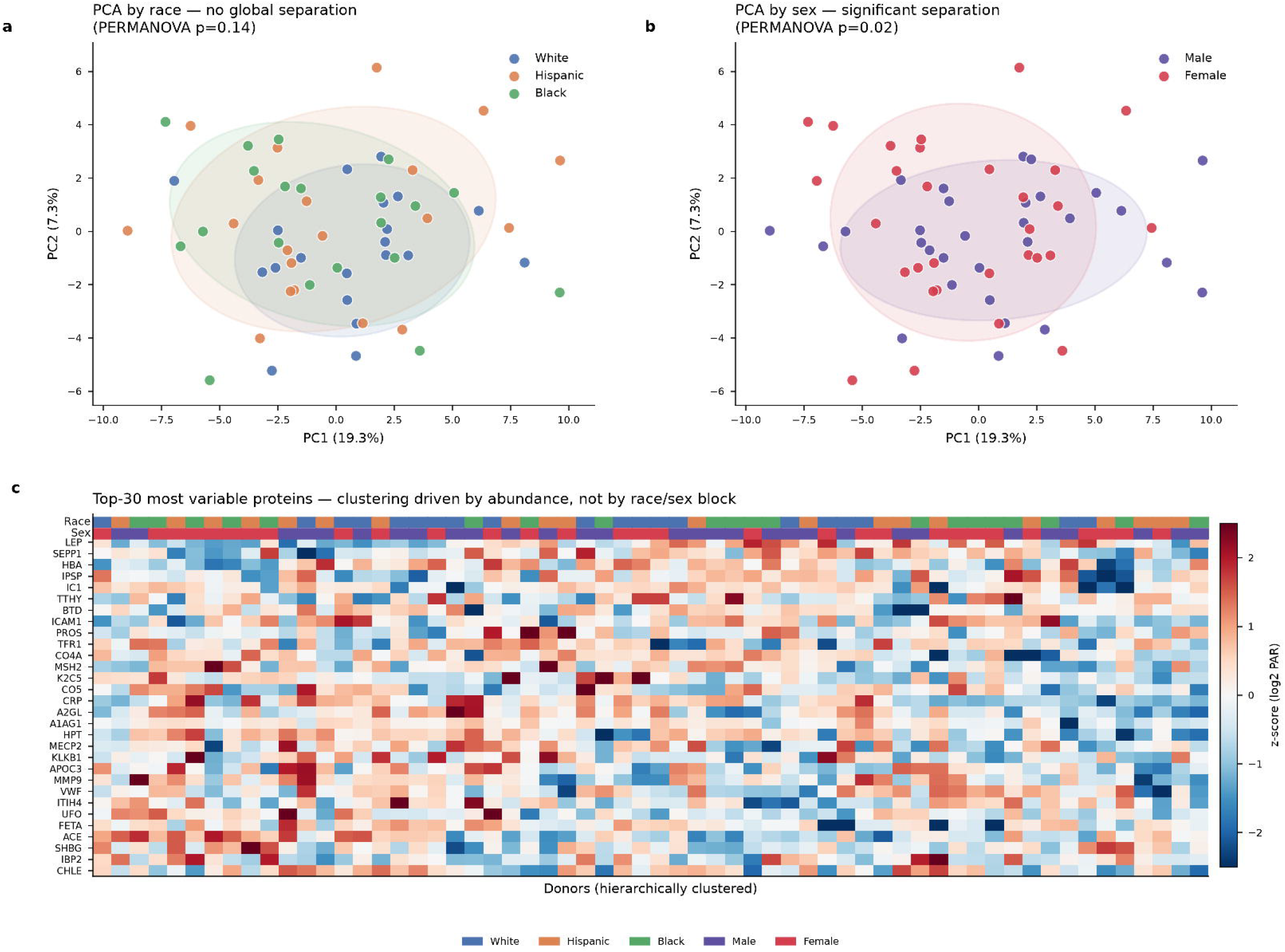
Global proteomic structure: sex is a systematic axis of variation; race is protein-specific. (a) Principal-component analysis (PCA) coloured by race shows no global separation (PERMANOVA p = 0.14; ellipses are 95% confidence regions). (b) The same PCA coloured by sex shows significant separation (PERMANOVA p = 0.02). PC1 and PC2 explain 19.3% and 7.3% of the variance, respectively. (c) Unsupervised hierarchical clustering of the 30 most variable proteins; donor clustering is driven by overall abundance rather than by race or sex blocks. Top annotation bars indicate race and sex; colour encodes the z-scored log□ peak-area ratio (PAR).

Per-protein two-way ANOVA reinforced this picture and quantified it. At a panel-wide FDR of q < 0.05, three proteins differed by sex and none by race or by the race × sex interaction; at a nominal threshold of p < 0.05, 13 proteins showed a sex effect, 5 a race effect and 7 an interaction (**Table 2, Table S2**). The contrast is instructive: sex produces the strongest and most reproducible signal across the panel and survives stringent multiple-testing correction, whereas race differences, though genuine, are concentrated in particular proteins and do not reshape the proteome globally. This asymmetry— global for sex, local for race—recurs throughout the analyses below and directly shapes the diagnostic implications.

**Table 2.**

| Gene | Protein | Status | Category | p(Race) | q(Race) | $\eta^2$(Race) | p(Sex) | q(Sex) | $\eta^2$(Sex) | log2FC(M/F) | p(Race×Sex) | q(Race×Sex) |
| --- | --- | --- | --- | --- | --- | --- | --- | --- | --- | --- | --- | --- |
| SHBG | Sex hormone-binding globulin | FDA | Endocrine/Metabolic | 0.440 | 0.832 | 0.030 | 0.000 | 0.002 | 0.284 | -0.518 | 0.175 | 0.830 |
| HBA | Hemoglobin subunit alpha | FDA | Iron/Hematology | 0.004 | 0.206 | 0.182 | 0.000 | 0.007 | 0.235 | 1.425 | 0.024 | 0.344 |
| ALBU | Serum albumin | FDA | Liver/Nutrition | 0.591 | 0.926 | 0.019 | 0.001 | 0.036 | 0.177 | 0.127 | 0.948 | 0.956 |
| PAFA | Platelet-activating factor acetylhydrolase (Lp-PLA2) | FDA | Lipid/Cardiovascular | 0.283 | 0.684 | 0.046 | 0.005 | 0.112 | 0.136 | 0.459 | 0.595 | 0.934 |
| MET | Hepatocyte growth factor receptor (c-Met) | LDT | Oncology | 0.780 | 0.943 | 0.009 | 0.009 | 0.156 | 0.120 | -0.317 | 0.204 | 0.830 |
| HEMO | Hemopexin | LDT | Iron/Hematology | 0.014 | 0.415 | 0.146 | 0.011 | 0.156 | 0.114 | -0.254 | 0.211 | 0.830 |
| RET4 | Retinol-binding protein 4 | FDA | Renal | 0.611 | 0.932 | 0.018 | 0.018 | 0.205 | 0.100 | 0.317 | 0.697 | 0.934 |
| IPSP | Plasma serine protease inhibitor (Protein C inhibitor) | LDT | Coagulation/Platelet | 0.787 | 0.943 | 0.009 | 0.022 | 0.205 | 0.093 | 0.507 | 0.156 | 0.830 |
| IBP3 | Insulin-like growth factor-binding protein 3 | FDA | Endocrine/Metabolic | 0.058 | 0.527 | 0.100 | 0.023 | 0.205 | 0.093 | 0.133 | 0.926 | 0.956 |
| APOC3 | Apolipoprotein C-III | LDT | Lipid/Cardiovascular | 0.988 | 0.988 | 0.000 | 0.024 | 0.205 | 0.091 | 0.316 | 0.617 | 0.934 |
| C1R | Complement C1r | LDT | Complement | 0.629 | 0.938 | 0.017 | 0.043 | 0.334 | 0.074 | -0.004 | 0.180 | 0.830 |
| FHR5 | Complement Factor H-related protein 5 | LDT | Complement | 0.503 | 0.854 | 0.025 | 0.048 | 0.334 | 0.071 | 0.359 | 0.714 | 0.934 |
| APOA2 | Apolipoprotein A-II | LDT | Lipid/Cardiovascular | 0.021 | 0.458 | 0.133 | 0.050 | 0.334 | 0.069 | -0.051 | 0.011 | 0.321 |
| TTHY | Transthyretin (prealbumin) | FDA | Liver/Nutrition | 0.036 | 0.527 | 0.116 | 0.094 | 0.452 | 0.051 | -0.389 | 0.016 | 0.344 |
| EGLN | Extracellular glycoprotein lactadherin (MFGE8) | LDT | ECM/Other | 0.117 | 0.527 | 0.076 | 0.129 | 0.455 | 0.042 | -0.088 | 0.011 | 0.321 |
| THRB | Prothrombin (Factor II) | FDA | Coagulation/Platelet | 0.202 | 0.566 | 0.058 | 0.392 | 0.742 | 0.014 | 0.072 | 0.043 | 0.537 |
| CD14 | Monocyte differentiation antigen CD14 (sCD14) | LDT | Inflammation/Immune | 0.005 | 0.206 | 0.180 | 0.611 | 0.868 | 0.005 | 0.111 | 0.887 | 0.952 |
| FIBA | Fibrinogen alpha | FDA | Coagulation/Platelet | 0.172 | 0.553 | 0.063 | 0.614 | 0.868 | 0.005 | -0.027 | 0.021 | 0.344 |
| MMP1 | Matrix metalloproteinase-1 | LDT | Oncology | 0.732 | 0.943 | 0.011 | 0.892 | 0.958 | 0.000 | -0.064 | 0.011 | 0.321 |

### Sex-associated proteins and their consistency across races

The FDR-significant sex effects were biologically coherent and, reassuringly, matched textbook physiology. Sex hormone-binding globulin (SHBG) was higher in females (log□ fold-change male/female = −0.52; partial η^2^ = 0.28; q = 0.0021), haemoglobin subunit α (HBA) was higher in males (log□FC = +1.43; η^2^ = 0.23; q = 0.0067), and serum albumin (ALBU) showed a smaller but significant sex difference (q = 0.036); all three are analytes of FDA-cleared assays (**Figure 3a,b**). The directions recapitulate well-established human biology—higher SHBG in women under oestrogenic regulation, and higher red-cell mass and haemoglobin in men—providing an internal biological validation of both the assay and the cohort.

**Figure 3.**
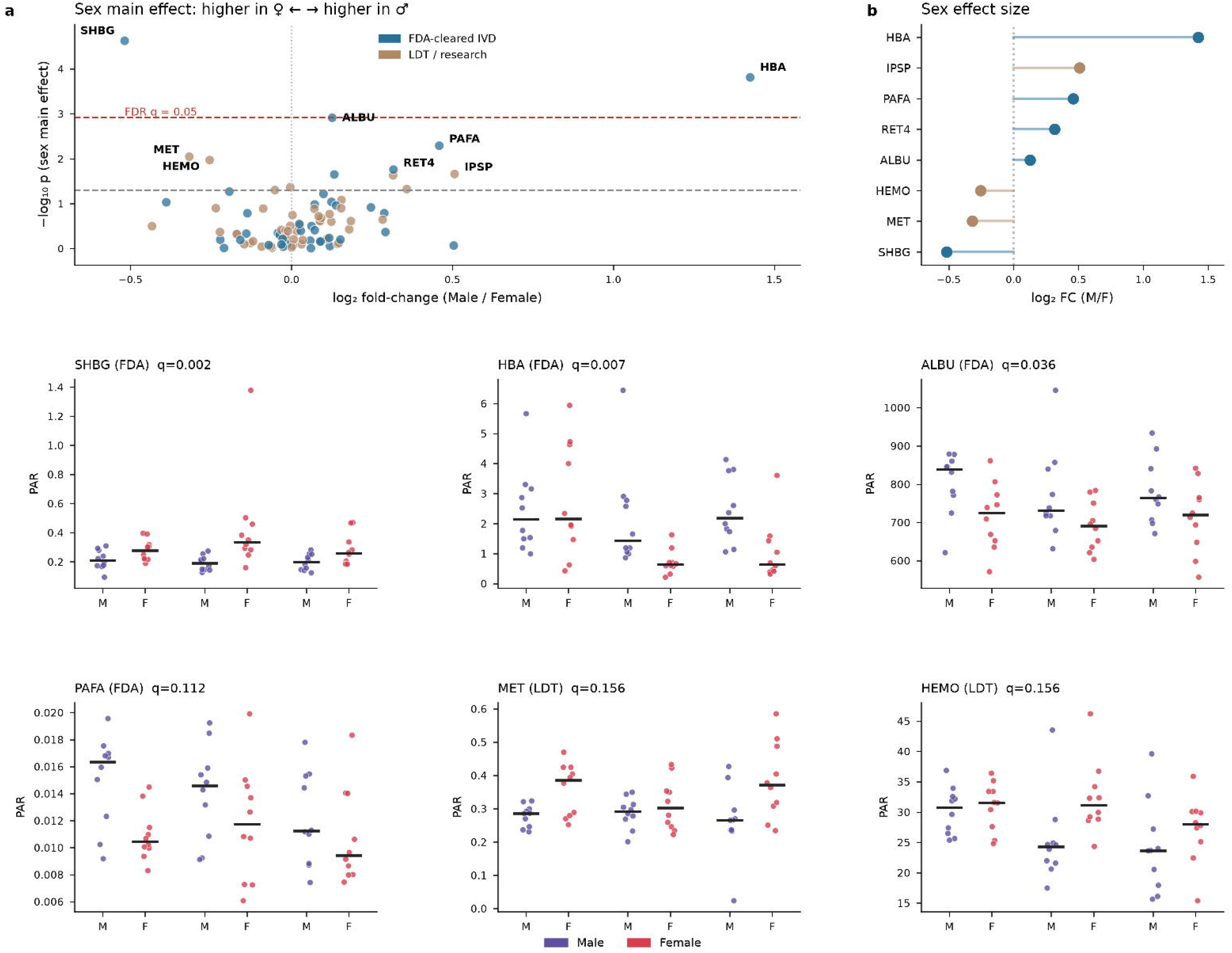
Race-associated differences in serum clinical-biomarker proteins. (a) Race main effect across 87 proteins (−log□□ p versus partial η^2^); points are coloured by regulatory status and the dashed line marks p = 0.05. No protein reached panel-wide FDR significance, but several showed nominal race effects (HBA, CD14, HEMO, APOA2, TTHY, A2GL). (b) Number of proteins with significant pairwise race contrasts (Tukey q < 0.05); most differences involve the Black-versus-White comparison. (c) Per-donor distributions for the leading race-associated proteins (PAR; horizontal bar = median; n = 20 per race). W = White, H = Hispanic, B = Black.

Critically for diagnostic use, these sex effects were not confined to a single race. When the male/female fold-change was computed separately within each race, the sex-associated proteins retained their direction across White, Hispanic and Black donors (**Figure 3c, Figure S2**); SHBG and HBA in particular pointed the same way in every group. The sex signature is therefore a stable, cross-population property of the healthy serum proteome rather than an artefact of any one demographic stratum—an important prerequisite if sex-specific reference intervals are to be transportable across diverse patient populations.

### Race differences are real but protein-specific

Although race did not restructure the proteome globally, individual proteins differed by race in reproducible, interpretable ways. Post-hoc Tukey testing identified five proteins with significant pairwise race contrasts (adjusted p < 0.05; **Figure 4**), most of them in the Black-versus-White comparison (**Figure 4a,b**). These comprised haemoglobin-α, transthyretin (TTHY), leucine-rich α-2-glycoprotein (A2GL), haemopexin (HEMO) and soluble CD14. Notably, soluble CD14 was higher in White (European-ancestry) than in Black donors (Figure 3c, CD14 panel), consistent with the well-documented enrichment of sCD14-raising CD14-locus alleles in European relative to African-ancestry populations;^7^ this recapitulation of a known ancestry-associated difference further supports the validity of the measurements.

**Figure 4.**
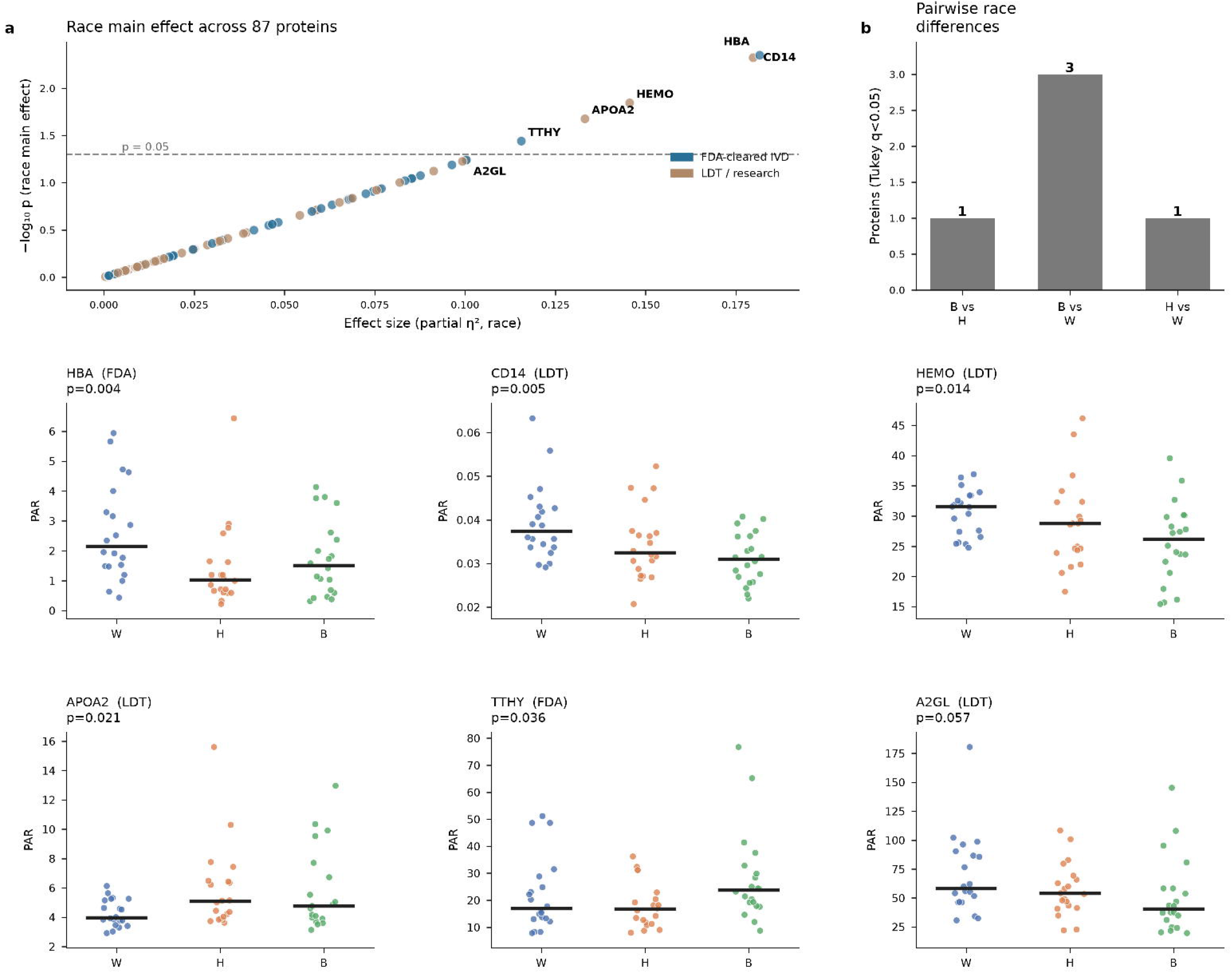
Sex-associated differences dominate serum biomarker variation and persist across races. (a) Sex main effect (−log□□ p versus log□fold-change male/female); the dashed grey line marks nominal p = 0.05 and the red line marks FDR q = 0.05. SHBG (higher in females) and HBA (higher in males) are the strongest signals. (b) Sex effect sizes (log□fold-change male/female) for the top sex-associated proteins. (c) Per-donor PAR distributions within each race block (White | Hispanic | Black); M = male (purple), F = female (red); bar = median (n = 10 per group). Sex directions are preserved across all three races.

We interpret these contrasts cautiously. They are best read as protein-specific, partly ancestry-linked differences in baseline abundance, not as evidence that “race” is a coherent biological variable. Where a difference can be traced to allele frequencies at the encoding gene, as for CD14, it is genetic ancestry—not the social category of race—that carries the signal, a distinction with direct bearing on whether and how demographic information should enter a reference interval.

### Demographic-agnostic reference intervals systematically misclassify subgroups

To translate these differences into diagnostic terms, we simulated the current default practice: a single reference interval applied to everyone. For each FDA-target protein carrying any demographic signal, we derived a pooled 95% reference interval from all 60 donors and asked what fraction of each race– sex subgroup fell outside it. A well-calibrated, unbiased interval flags 5% of any healthy subgroup by definition. Instead, several subgroups were flagged far more often, reaching ∼25% for individual proteins (for example, SHBG in Hispanic females and biotinidase in Black males; **Figure 5a,b**). The burden was unevenly distributed: Hispanic females carried the heaviest cumulative load, exceeding the 10% misclassification level for eight FDA targets, and the disparity was driven by both sex and race strata (**Figure 5c,d**).

**Figure 5.**
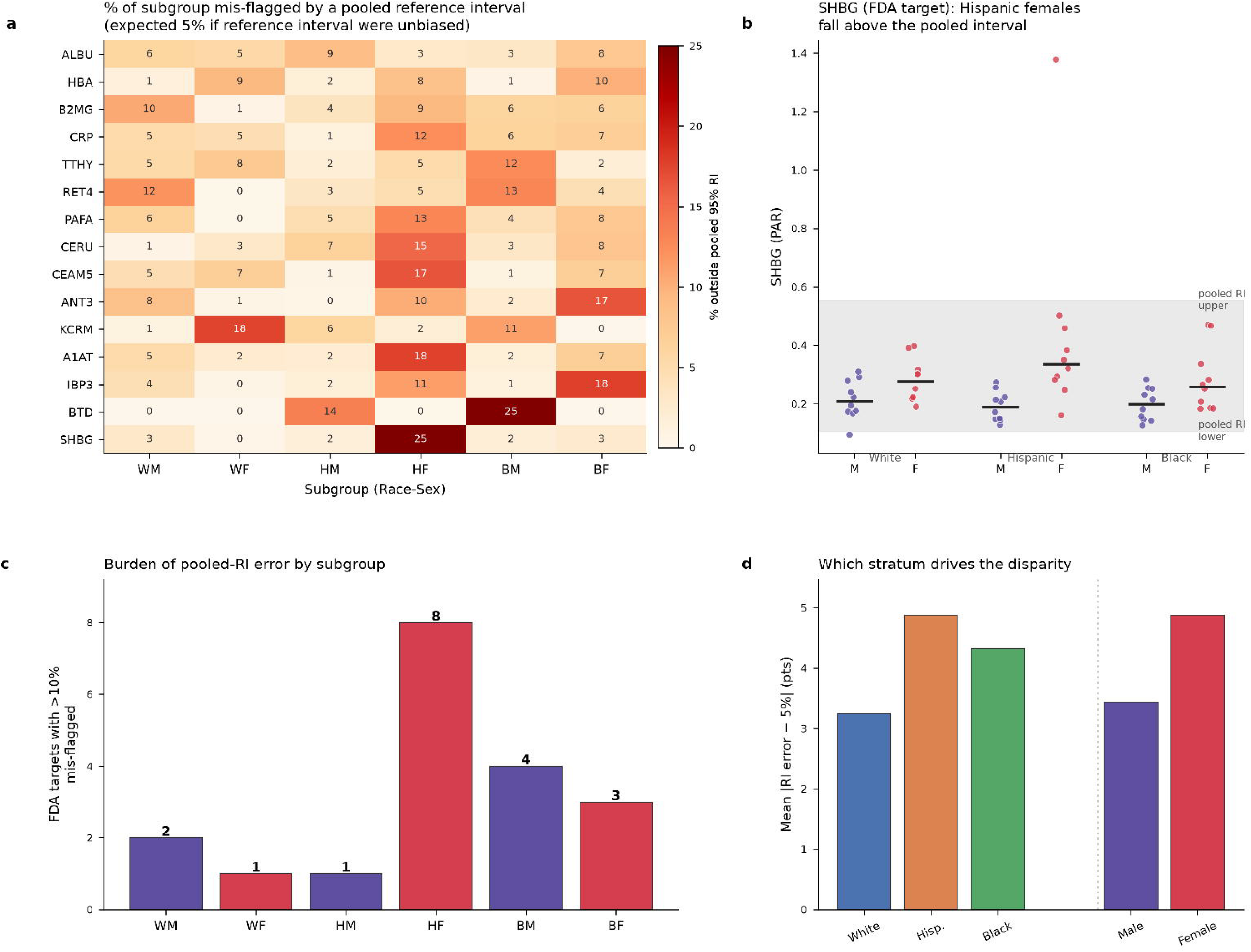
Clinical impact: race/sex-agnostic reference intervals systematically misclassify subgroups. (a) Percentage of each race–sex subgroup falling outside a single pooled 95% reference interval, by protein; an unbiased interval would flag 5%. (b) SHBG (an FDA target): Hispanic females fall disproportionately above the pooled interval (shaded band = pooled reference interval; points = donors; bars = subgroup medians). (c) Cumulative burden: number of FDA targets for which each subgroup is mis-flagged at >10%; Hispanic females carry the heaviest burden. (d) Mean absolute deviation from the expected 5% mis-flagging rate, decomposed by race and by sex stratum. Subgroups (WM, WF, HM, HF, BM, BF) are as defined in Figure 1.

Because every donor here is healthy, each such flag is a false positive attributable solely to demographic mismatch between the individual and the pooled reference population. The error is not random noise that averages out across a population; it is systematic and direction-consistent, concentrating in particular subgroups and particular proteins. For a marker used to rule disease in or out, a bias of this magnitude at the reference-interval boundary translates directly into differential false-positive—and, symmetrically, false-negative—rates across demographic groups.

## Discussion

Across a broad, single-platform panel of clinically used biomarker proteins measured in demographically balanced healthy donors, baseline serum concentrations carry a reproducible sex signature and protein-specific race differences. The sex signal is the more pervasive: it is visible in unsupervised multivariate structure, anchored by physiologically expected markers (SHBG, haemoglobin), survives stringent multiple-testing correction, and is consistent in direction across all three races. Race differences are genuine but concentrated in particular proteins—CD14, transthyretin, haemoglobin, A2GL and haemopexin—rather than reshaping the proteome as a whole. This global-versus-local asymmetry is itself a substantive finding, and it aligns with large affinity-based proteomic surveys in which sex exerts broad effects across the circulating proteome.^17, 19^

The practical consequence is concrete and quantifiable. When a single, demographic-agnostic reference interval is applied—the prevailing default for many protein assays—healthy individuals from specific subgroups are mis-flagged as abnormal at rates well above the nominal 5%, approaching one in four for some FDA-target proteins in specific race–sex groups. Because the affected proteins include analytes of cleared IVDs, this is not a hypothetical concern confined to research markers. A direction-consistent bias at the decision boundary erodes both diagnostic accuracy and equity, and it does so silently, because the flagged individuals are in fact healthy.

Mechanistically, the sex effects we observe are readily understood: SHBG is oestrogen-regulated and higher in women, whereas higher haemoglobin in men reflects androgen-driven erythropoiesis. Their reproducibility across races makes sex-specific intervals an obvious, low-risk refinement, and sex partitioning is already standard for a handful of analytes.^3^ Ancestry is more delicate. Where a race-associated difference is traceable to allele frequencies at the encoding locus—CD14 being the clearest case^7^—the biologically meaningful variable is genetic ancestry, not the social category of race. This distinction matters, because the past decade has shown how uncritically embedding race in clinical algorithms can entrench inequity; the corrective is not to ignore ancestry-linked biology but to measure it directly and represent it appropriately.^8, 10, 29, 30^

These findings argue that demographic context belongs in the interpretation of protein biomarkers— through sex- and, where mechanistically warranted, ancestry-informed reference intervals and decision thresholds, rather than a single pooled interval. Targeted MRM is well suited to establishing such intervals at scale: its internal standardization makes measurements portable across laboratories and over time, the property that has made it a reference approach for clinical biomarker verification.^23,25,26^ As multiplexed proteomic risk models move toward the clinic, building demographic calibration in from the outset—rather than retrofitting it after disparities emerge—would help ensure that the benefits of proteomic diagnostics are shared equitably.^21^

Our study has limitations. The cohort comprises healthy young adults (20–30 y) across three self-reported race categories, sampled as commercial single-donor sera with n = 10 per race–sex group; the design is balanced and age-matched but modest in size and narrow in age range, and self-reported race captures genetic ancestry only imperfectly. The reference-interval simulation is illustrative of the direction and approximate magnitude of the effect rather than a clinical validation, and quantitation is relative (peak-area ratio to SIS peptides) rather than absolute concentration. Larger, absolutely quantified and more demographically granular cohorts—spanning the full adult age range and, ideally, using measured genetic ancestry rather than self-reported race—will be needed to derive deployable, partitioned reference intervals. That said, the concordance of the sex effects with established physiology and of specific race effects (notably CD14) with prior genetic literature gives us confidence in the core conclusion.

In sum, demographic-agnostic reference intervals introduce a predictable, direction-consistent bias into the interpretation of clinically used protein biomarkers. Incorporating demographic context into reference intervals and decision thresholds is therefore not a concession to complexity but a step toward more accurate and more equitable protein-based diagnostics.

## Data and Code availability

The mass spectrometry data have been deposited to Panorama Public (https://panoramaweb.org/), and are accessible at https://panoramaweb.org/9KcVFW.url. All analysis scripts used in this study, including cosine similarity calculation, differential abundance analysis (limma), and figure generation, are publicly available on GitHub (https://github.com/kimlab-cnu/ProteoBias). Additional data are available from the corresponding author on reasonable request.

## Acknowledgement

We acknowledge the use of Claude Opus 4.8 solely for linguistic refinement and grammatical corrections in manuscript preparation. All scientific content, data analysis, and intellectual contributions presented herein were developed independently by the authors without the use of generative AI tools.

## Author Contributions

A.S. and H.K.: conceptualization, methodology, experimental analysis, data analysis, visualization; J.J., E.H., Y.C., J.P., H.L., and S.P.: data analysis; A.S, and H.K.: writing-original draft, conceptualization, project administration, resources, supervision, writing-review & editing. All authors have read and approved the final manuscript.

