## Supplementary figures and images for "Sex and race shape serum protein biomarkers and bias diagnostic reference intervals"

### Figure S1

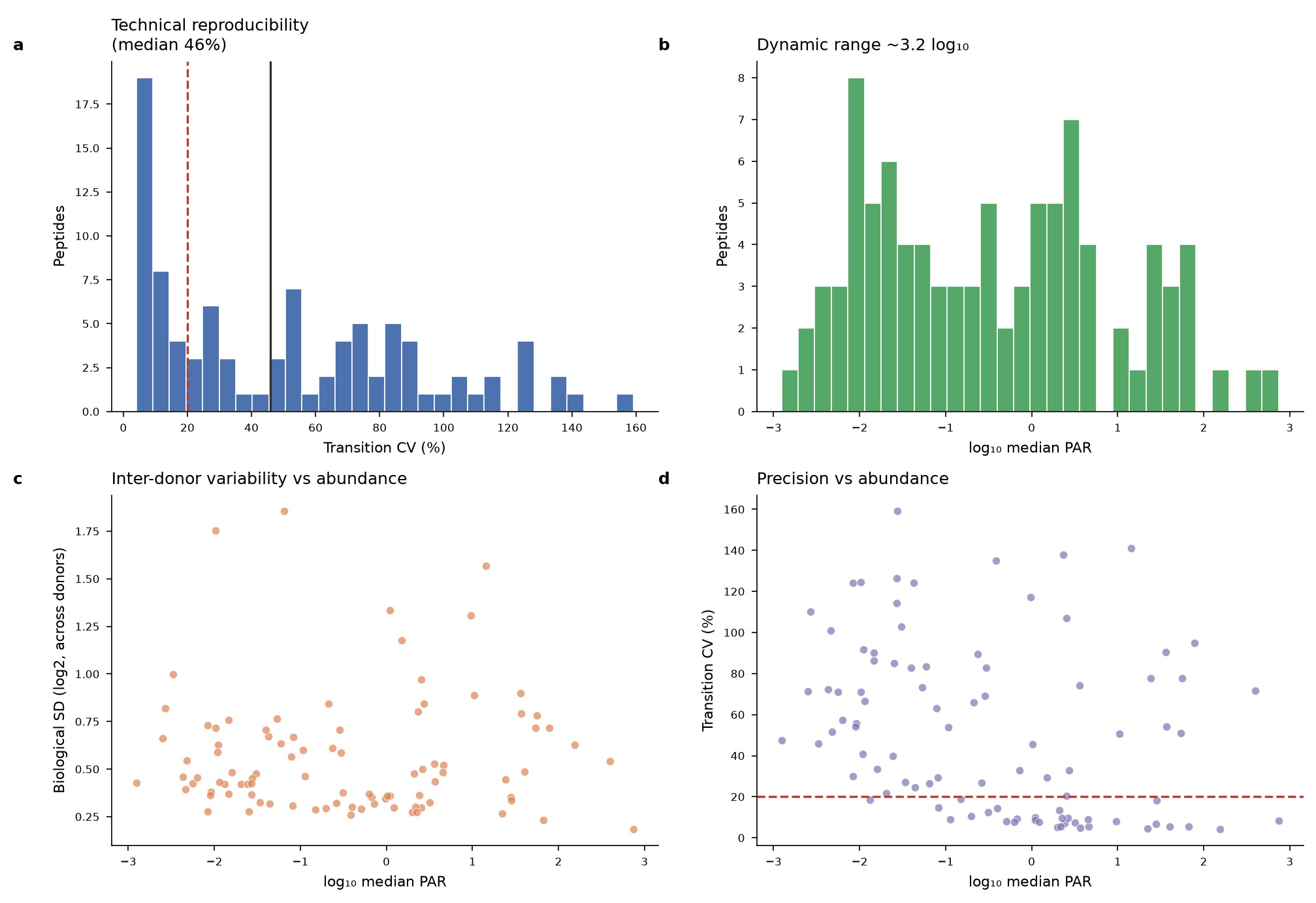

### Figure S2

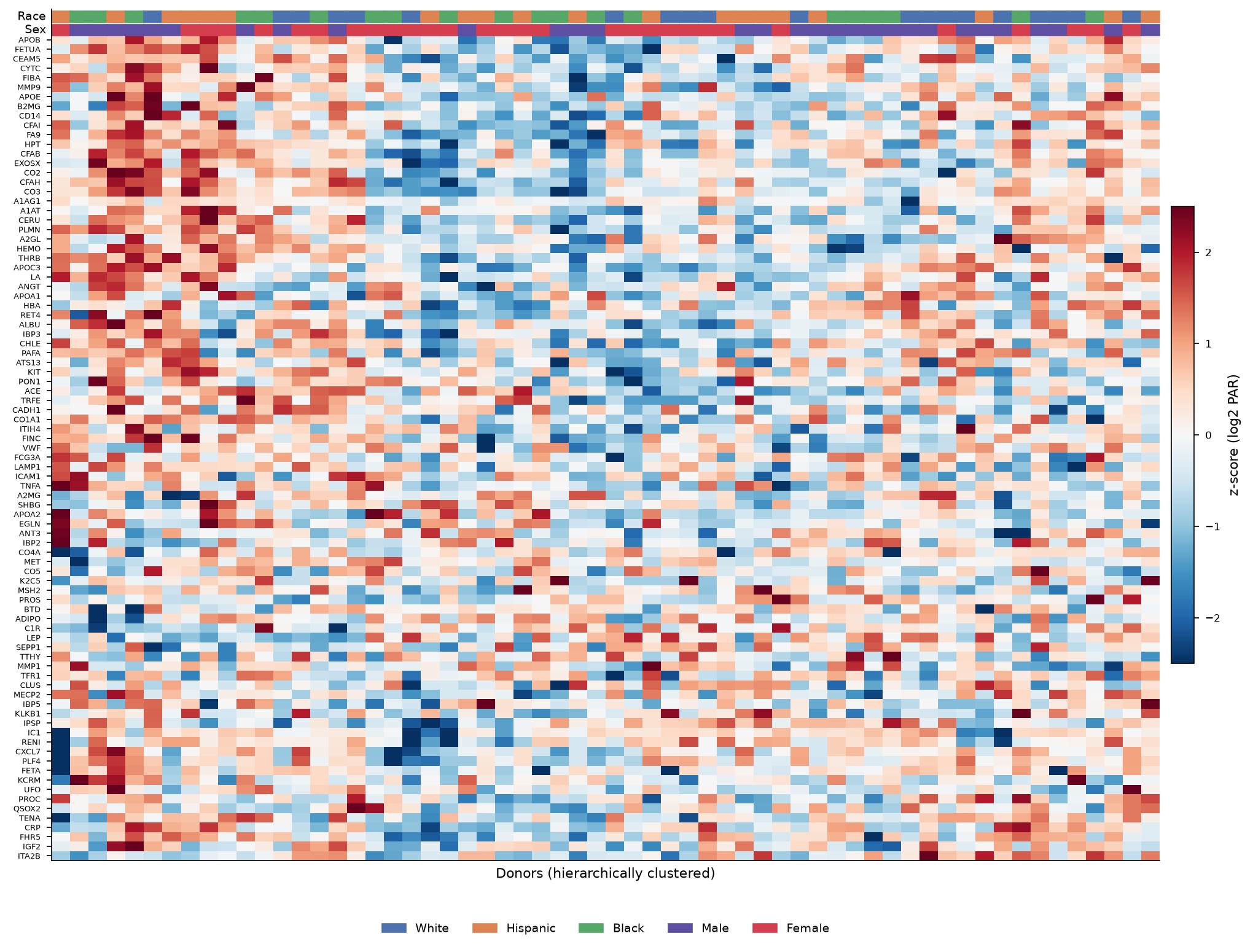

### Figure S3

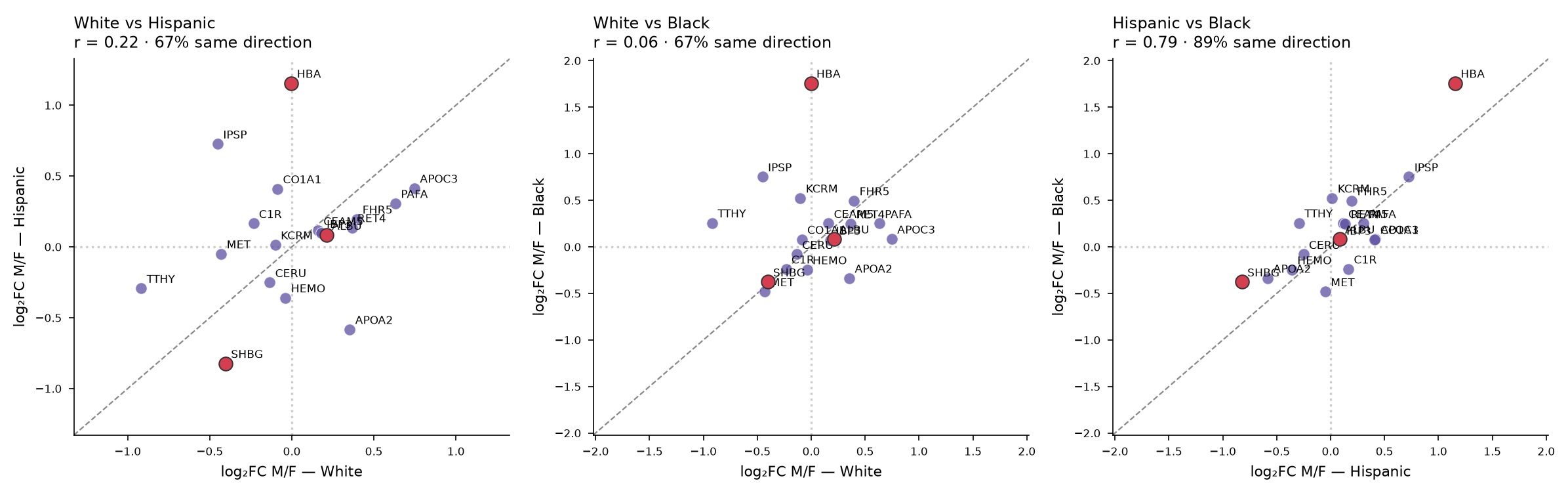
