## Supplemental Text for "Sex and race shape serum protein biomarkers and bias diagnostic reference intervals"

**Supplementary Figures**

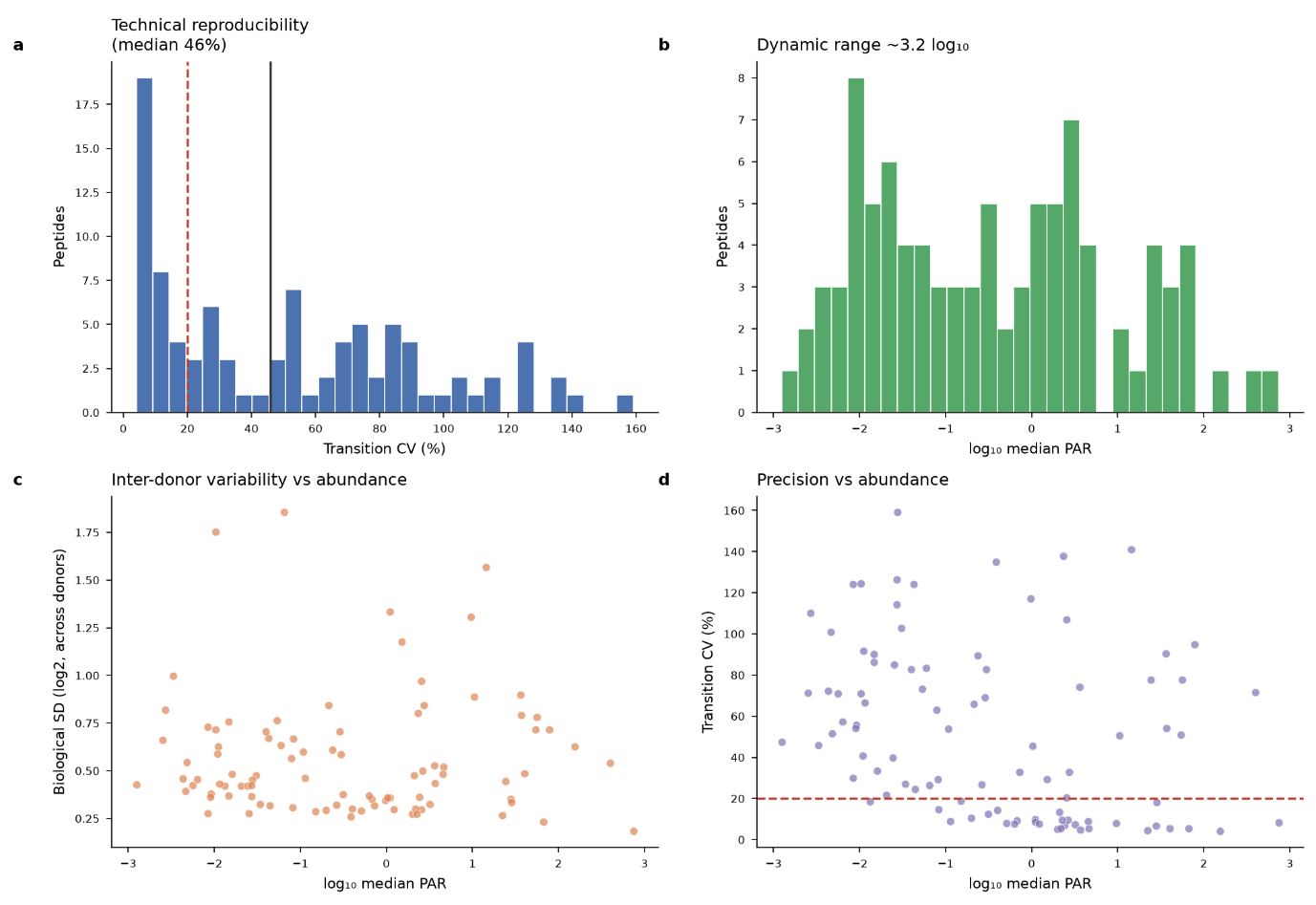

**Figure S1. Assay quality control.**

(a) Distribution of inter-transition coefficients of variation across the 93 peptides (median ≈ 41%; 20% reference line). (b) Dynamic range of median PAR across ~6 orders of magnitude. (c) Inter-donor biological variability versus abundance. (d) Analytical precision (transition CV) versus abundance, showing the expected loss of precision for the lowest-abundance targets.

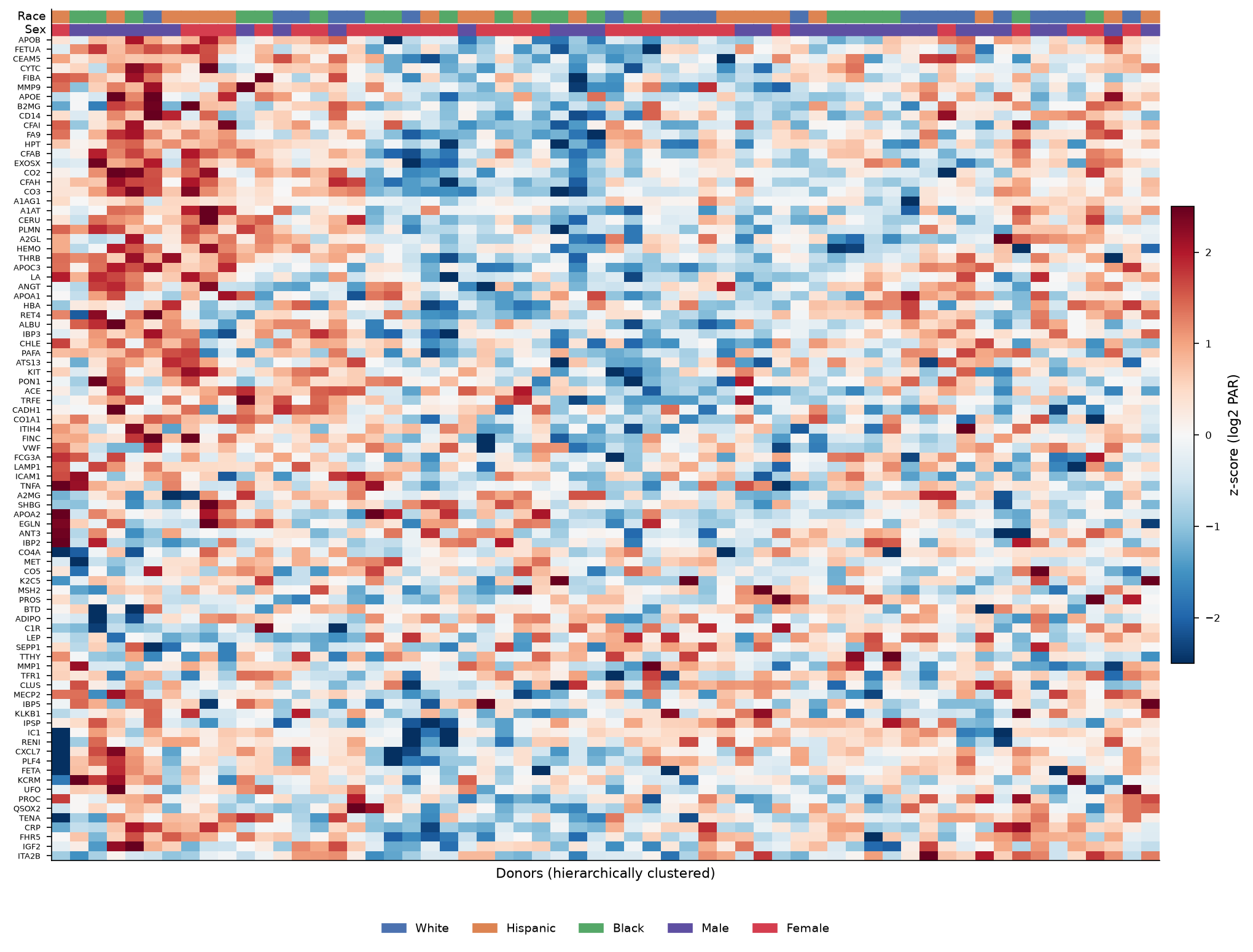

**Figure S2. Complete clustered heatmap.**

Z-scored log₂ PAR for all 87 proteins across all 60 donors, with Ward-linkage hierarchical clustering of both proteins and donors and race/sex annotation strips. Donor clustering is governed by overall abundance structure rather than by race or sex block membership.

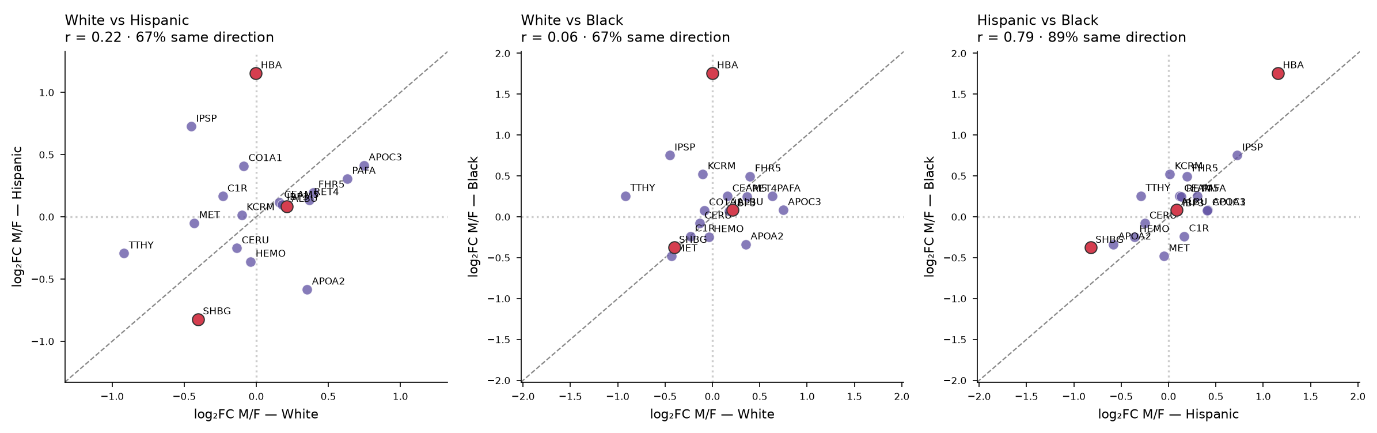

**Figure S3. Cross-race consistency of the sex effect.**

Pairwise comparison of within-race male/female log₂ fold-changes for the sex-associated proteins (nominal p < 0.1). Points on the diagonal indicate identical sex effects between two races; the percentage of proteins sharing the same direction of effect is annotated. The strongest markers (SHBG, HBA) are concordant across all three race pairings, demonstrating that the sex signature is a stable, cross-population property.

**Supplementary Tables**

Table S1. Full target annotation (87 proteins): gene symbol, UniProt accession, protein name, FDA/LDT clinical status, detailed and broad clinical category, and clinical use.

Table S2. Reference-interval misclassification: percentage of each race–sex subgroup expected to fall outside the pooled 95% reference interval, per analyzed FDA-target protein.

Table S3. Complete two-way ANOVA results (87 proteins): p-values, FDR q-values and partial η² for race, sex and race × sex effects, plus male/female log₂ fold-change.

Table S4. Group median PAR for all 87 proteins across the six race–sex groups.

Table S5. Peptide-level QC summary: per-peptide median PAR, transition CV, inter-donor SD and dynamic range.
